## Supplementary Material for "Negative feedback may suppress variation to improve collective foraging performance"

### Supplementary text of the article
### Negative feedback may suppress variation to improve collective foraging performance

Andreagiovanni Reina and James A. R. Marshall

January 12, 2022

#### ST1. Relationship between transition rates and food patch quality

Through numerical optimisation, we defined for both models the functional form of the transition rates that approximates best the target distribution. The optimisation routine computes the cumulative sum of errors of the mean-field model. The error is summed for all $n$ food patches. This sum is also cumulative with respect to the time dimension, that is, the sum of errors is computed every time unit until the end of the simulation at $T_{\max}$, as:

$$\Theta = \sum_{t=1}^{T_{\max}} \sum_{i=1}^{n} \left| x_i^t - \frac{q_i}{\sum_{j \in n} q_j} \right| \;, \tag{SE1}$$

where $|\,.\,|$ is the operator for absolute value. The optimisation routine explores the parameter space to find the values that minimise $\Theta$ of Eq. (SE1). In order to find parameters that would work well for different numbers of food patches $n$, different quality values, and for different initial conditions, we compute $\Theta$ (the cumulative sum of errors of Eq. (SE1)) for several setups, and use their sum as the value to minimise. We test both $n = 2$ and $n = 3$. For $n = 2$, we fix quality $q_2 = 0.5$ and compute $\Theta$ for $q_1 \in \{0, 0.1, 0.2, ..., 1.0\}$. For $n = 3$, we fix qualities $q_2 = 0.6$ and $q_2 = 0.3$, and compute $\Theta$ for $q_1 \in \{0, 0.1, 0.2, ..., 1.0\}$. We also test all possible initial states with sub-populations committed to the two, or three, options with step 0.1 (e.g., $(x_1, x_2, x_3) \in \{(0,0,0),(0,0,0.1),(0,0,0.2),...,(0.2,0.6,0.1),...,(1,0,0)\}$). The sum of $\Theta$ in all these setups is the quantity that is minimised by the optimisation routine. The optimisation code and results are available on GitHub at `https://github.com/DiODeProject/VarianceSuppression`.

We set the rates as logistic functions of the food patch quality,

$$\lambda = \frac{2\bar{\lambda}}{e^{-s(q_i - 0.5)} + 1} \;, \tag{SE2}$$

where $\lambda$ is a generic rate, $\bar{\lambda}$ is the average rate frequency, and $s$ determines the slope of the curve[1]. Figure SF1 shows some representative curves of Eq. (SE2) for different values of the slope $s$. The optimisation routine selects values in the range $s \in [0, 5]$, representing at the two extremes the constant function ($s = 0$, quality-insensitive), and approximately a linear relationship ($s = 5$, $\lambda \approx 2\bar{\lambda} q_i$).

We run the optimisation routine in a variety of configurations to determine both the best relative strength between the transition rates (rate strength $\bar{\lambda} \in (0, 1000]$ for abandonment, recruitment, and stop signalling) and their functional form (slope $s$ for all rates). The process speed is always normalised on the average rate of discovery, which is fixed at value $\bar{\lambda} = 1$ in every test. The results are reported in Table 1 to 6 for various configurations. These results motivated the definition of the two streamlined models that we presented in the main text, as indicated in the following.

**Strong recruitment is most beneficial when combined with stop signalling.** In Table 1, we allowed the optimisation routine to freely select the best average rate strengths and slope for all rates. Instead in Table 2, we run the same process but fixed stop-signalling strength to zero (positive

[1] Note that the abandonment rate can be inversely proportional to the food patch quality, therefore the term $q_i$ is replaced by $(1 - q_i)$.

| $T_{\max}$ | Discovery | | Recruitment | | Abandonment | | Stop signalling | | error $\Theta$ |
|---|---|---|---|---|---|---|---|---|---|
| | $\bar{\lambda}$ | slope $s$ | $\bar{\lambda}$ | slope $s$ | $\bar{\lambda}$ | slope $s$ | $\bar{\lambda}$ | slope $s$ | |
| 10 | 1 | 3.18753 | 993.266 | 3.55255 | 1.50558 | 1.64438 | 25.8788 | 0.00355132 | 0.167723 |
| 100 | 1 | 4.01337 | 990.897 | 3.59278 | 0.722135 | 1.57534 | 12.811 | 0. | 1.61001 |
| 1000 | 1 | 4.86449 | 973.061 | 3.66919 | 0.229161 | 1.86251 | 4.84565 | 0. | 15.5916 |

Table 1: Optimised parameters with all rates and slopes free (expect for discovery $\bar{\lambda} = 1$).

| $T_{\max}$ | Discovery | | Recruitment | | Abandonment | | Stop signalling | | error $\Theta$ |
|---|---|---|---|---|---|---|---|---|---|
| | $\bar{\lambda}$ | slope $s$ | $\bar{\lambda}$ | slope $s$ | $\bar{\lambda}$ | slope $s$ | $\bar{\lambda}$ | slope $s$ | |
| 10 | 1 | 4.0017 | 3.82747 | 1.97406 | 0.862282 | 0.00372217 | 0 | 0 | 0.71383 |
| 100 | 1 | 4.34572 | 0.776195 | 1.94331 | 0.166382 | 0.138089 | 0 | 0 | 3.60998 |
| 1000 | 1 | 4.09265 | 11.2862 | 0.0360765 | 0.294101 | 0.0811387 | 0 | 0 | 21.5086 |

Table 2: Optimised parameters with all rates and slopes free, except for discovery strength fixed to 1 and stop-signalling strength fixed to zero, i.e. model without negative feedback.

| $T_{\max}$ | Discovery | | Recruitment | | Abandonment | | Stop signalling | | error $\Theta$ |
|---|---|---|---|---|---|---|---|---|---|
| | $\bar{\lambda}$ | slope $s$ | $\bar{\lambda}$ | slope $s$ | $\bar{\lambda}$ | slope $s$ | $\bar{\lambda}$ | slope $s$ | |
| 10 | 1 | 4.50084 | 100. | 3.75795 | 0.481693 | 2.57493 | 8.09239 | 0.0495871 | 0.240978 |
| 100 | 1 | 4.85832 | 100. | 3.70402 | 0.206428 | 2.04744 | 3.8593 | 0.0131419 | 1.85758 |
| 1000 | 1 | 4.61453 | 100. | 3.56596 | 0.0947592 | 1.82524 | 1.3963 | 0.0672265 | 16.3936 |

Table 3: Optimised parameters with recruitment strength fixed to 100 and discovery strength $\bar{\lambda} = 1$. Values of all other rates and slopes are free.

| $T_{\max}$ | Discovery | | Recruitment | | Abandonment | | Stop signalling | | error $\Theta$ |
|---|---|---|---|---|---|---|---|---|---|
| | $\bar{\lambda}$ | slope $s$ | $\bar{\lambda}$ | slope $s$ | $\bar{\lambda}$ | slope $s$ | $\bar{\lambda}$ | slope $s$ | |
| 10 | 1 | 3.65107 | 100. | 0.0361097 | 9.99617 | 0.0463419 | 0 | 0 | 0.845349 |
| 100 | 1 | 4.04299 | 100. | 0.0268109 | 5.74431 | 0.00317431 | 0 | 0 | 3.93436 |
| 1000 | 1 | 4.02302 | 100. | 0.0108346 | 2.12533 | 0.00579289 | 0 | 0 | 21.825 |

Table 4: Optimised parameters with recruitment strength fixed to 100, stop-signalling strength fixed to zero, and discovery strength fixed to 1. Values of all other rates and slopes are free.

| $T_{\max}$ | Discovery | | Recruitment | | Abandonment | | Stop signalling | | error $\Theta$ |
|---|---|---|---|---|---|---|---|---|---|
| | $\bar{\lambda}$ | slope $s$ | $\bar{\lambda}$ | slope $s$ | $\bar{\lambda}$ | slope $s$ | $\bar{\lambda}$ | slope $s$ | |
| 10 | 1 | 5. | 1. | 5. | 0.171573 | 5. | 0.586042 | 0. | 0.619 |
| 100 | 1 | 4.97243 | 1. | 3.74725 | 0.076198 | 1.96719 | 0.176224 | 0.110512 | 3.19643 |
| 1000 | 1 | 4.42046 | 1. | 3.11191 | 0.0413467 | 0.879217 | 0.0486974 | 0.00521406 | 20.1137 |

Table 5: Optimised parameters with both recruitment and discovery strengths fixed to $\bar{\lambda} = 1$. Values of all other rates and slopes are free.

| $T_{\max}$ | Discovery | | Recruitment | | Abandonment | | Stop signalling | | error $\Theta$ |
|---|---|---|---|---|---|---|---|---|---|
| | $\bar{\lambda}$ | slope $s$ | $\bar{\lambda}$ | slope $s$ | $\bar{\lambda}$ | slope $s$ | $\bar{\lambda}$ | slope $s$ | |
| 10 | 1 | 4.53101 | 1. | 4.92961 | 0.500082 | 0.321492 | 0 | 0 | 0.719155 |
| 100 | 1 | 4.35149 | 1. | 1.82299 | 0.180247 | 0.0761111 | 0 | 0 | 3.61027 |
| 1000 | 1 | 4.09204 | 1. | 1.04416 | 0.0643803 | 0.117602 | 0 | 0 | 20.9055 |

Table 6: Optimised parameters with both recruitment and discovery strengths fixed to $\bar{\lambda} = 1$, and stop-signalling fixed to zero. Values of all other rates and slopes are free.

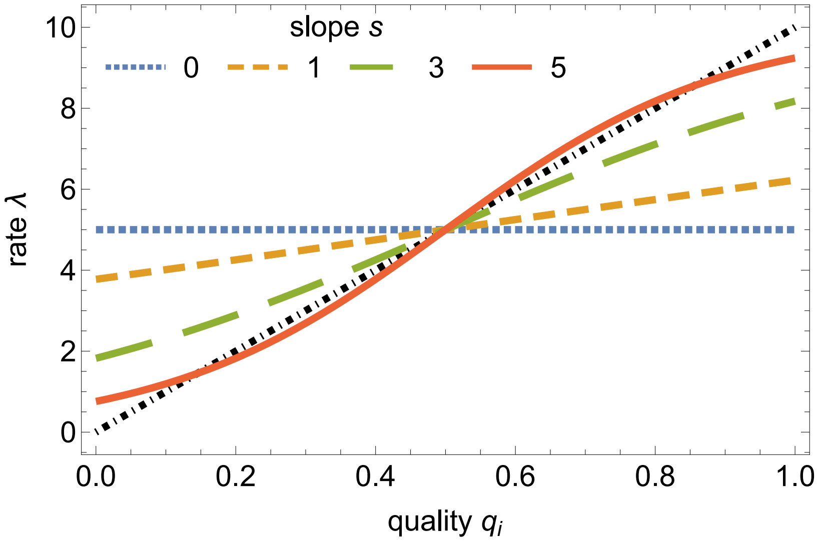


Figure SF1: The relationship between the food patch quality $q_i$ (x-axis) and the transition rate $\lambda$ (y-axis) is modelled with Eq. (SE2). We report four representative examples for $\bar{\lambda} = 5$ and slope values $s \in \{0, 1, 3, 5\}$. With $s = 0$ (blue dotted line), the rate is quality-independent, *i.e.* $\lambda = \bar{\lambda}$; instead, with $s = 5$ (orange solid curve), the relationship between $\lambda$ and $q_i$ is approximately linear (the linear function $\lambda = 2\bar{\lambda}q_i$ is included for comparison as a black dot-dashed line).

feedback only). When stop signalling was possible, the optimised recruitment reached values close to the highest boundary ($\sim 1000$). Instead, the system with disabled negative feedback (*i.e.* stop signalling $\bar{\lambda} = 0$) set recruitment to values two to three orders of magnitude smaller than when inhibition is enabled. Comparing the errors resulting from the two configurations, for every tested $T_{\max}$ (*i.e.* quick or slow dynamics), the system with both positive and negative feedback always has a lower error $\Theta$.

**Recruitment is proportional to quality only when there is negative feedback.** In Table 3 and 4, we fixed the recruitment strength to a high value, 100, and optimised all other parameters for the system with and without negative feedback, respectively. We can see that in the system without negative feedback (Table 4), the optimised slope for recruitment has values close to 0, therefore recruitment is quality-insensitive. On the contrary, the system with negative feedback (Table 3) has high values of slope $s$, approximating a linear relationship. Therefore, in the two models of the main text, we selected the relationships that minimised the error and could best approximate the target distribution: we set the recruitment rate in the model with positive feedback only as constant, and in the model with negative feedback as linearly proportional to the quality $q_i$.

**Stop signalling is relatively small and constant.** In Tables 1, 3, and 5, we can see that stop signalling strength $\bar{\lambda}$ is always one to two orders of magnitude smaller than recruitment and, most importantly, always has a slope close to zero. A slope close to zero means that the best results are given when stop signalling is constant and independent of the quality value. Therefore, in the main text, the model has constant stop signalling, with strength optimised to the recruitment strength.

#### ST2. Parameters

In our study, the quality $q_i$ is normalised in the range $q_i \in [0, 1]$. The abandonment rate is set to a constant low value $a = 10^{-3}$. The two models have different recruitment rates $r_i$: the model with negative feedback scales linearly with the option's quality, *i.e.* $r_i = \rho q_i$, and the model without negative feedback has a quality-independent rate $r_i = \rho$. The scaling factor $\rho$ that tunes the recruitment strength is set in order to have the same average rate $r$ in both models. Therefore, assuming that the expected quality $E(q)$ is the average of the range $[0, 1]$, thus $E(q) = 0.5$, we have that $\rho$ in the model without negative feedback is half of $\rho$ in the model with negative feedback. In every

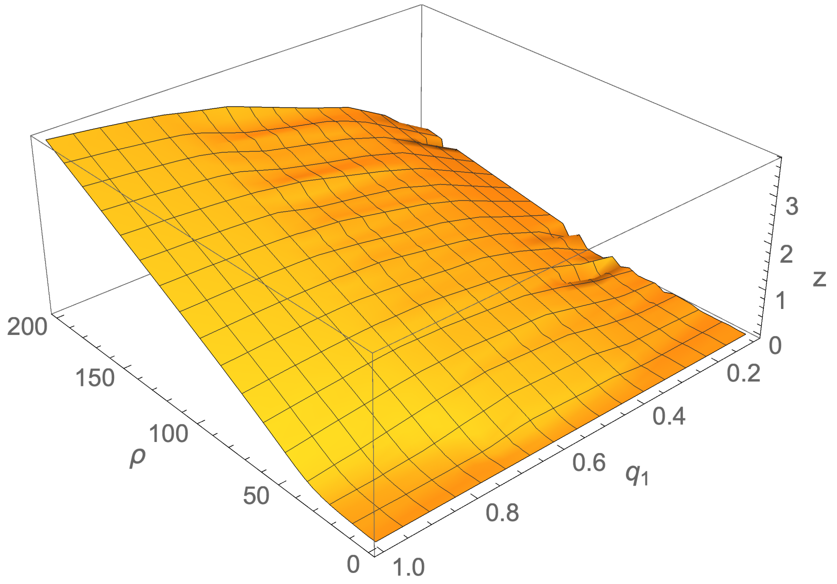


Figure SF2: The negative social feedback is numerically computed for each tested value of positive social feedback strength $\rho$ and food patch quality $q_1$ (for two-patches environments and $q_2 = 0.5$). The optimal negative social feedback is linearly proportional to $\rho$ and $q_1$.

plot of this paper, we indicate $r$, the average value of the possible rates $r_i$. The self-inhibition strength $z$ is tuned through numerical simulation to the best value in terms of speed and accuracy. In particular, we computed the strength of $z$ which took the system to a sum of squared error $SSE < 10^{-4}$. The squared error is computed as the sum of the squared distance from the target distribution for every population. Figure SF2 shows that the best stop signalling strength increases with the positive social feedback $\rho$. Figure 3 of the main text shows how the stop signalling strength influences the speed and the robustness of the system. As documented in previous work [1, 2], stronger signalling leads to quicker but less robust, or less accurate, dynamics.

#### ST3. Stability analysis

The dynamics of the two systems, described in the main text through chemical reactions (see Table 1 in the main text), can also be described via ODE systems. The system without negative social feedback reads as:

$$\begin{aligned} \dot{x}_i &= q_i\, x_U + x_i(\rho\, x_U - a), \qquad i \in \{1, \ldots, n\} \\ x_U &= 1 - \textstyle\sum_{i=1}^{n} x_i \;, \end{aligned} \tag{SE3}$$

where $x_i$ is repeated for the $n$ subpopulations committed to the $n$ patches. The system of Eq. (SE3) has two fixed points. Only one of the points is stable and exists in the solution domain relevant to our study, *i.e.* within $0 \leq x_i^* \leq 1$. This point is

$$x_i^* = \frac{q_i}{\sum_{j=1}^{n} q_j}\, \epsilon \,, \tag{SE4}$$

where $\epsilon \approx 1$, for small leak $a \ll 1$. Therefore, the expected solution accurately approximates the target distribution of Eq. (1) in the main text.

The $n$ eigenvalues of the system of Eq. (SE3) at the stable fixed point of Eq. (SE4) are all negative. However, one of the eigenvalues has the absolute value several order of magnitude larger than the value of the other $n-1$ eigenvalues which are approximately equal to zero. This asymmetry in the eigenvalue strength explains the type of dynamics observed in the system (*e.g.* Figure 1 bottom-left of the main text). The population quickly gets committed to the $n$ food patches, *i.e.* with $x_u \approx 0$, and then imperceptibly slowly moves towards the point of Eq. (SE4) on the $(n-1)$-dimension plane $x_1 = x_2 = \cdots = x_n$. However, the attraction is so low that practically the system does not move once it reaches that plane.

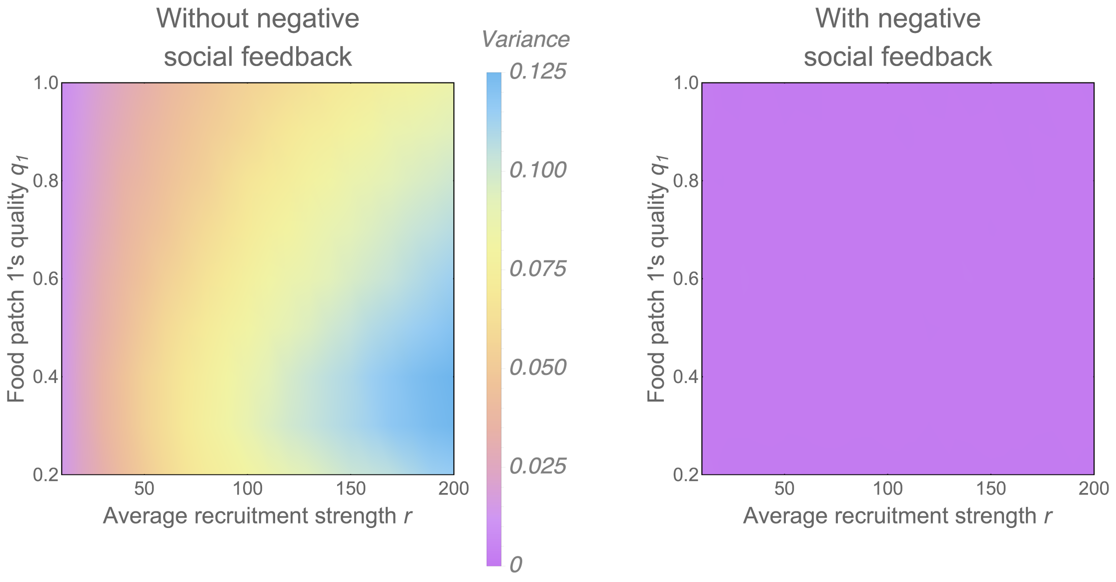


Figure SF3: Variance of the model without (left) and with (right) negative social feedback for varying average recruitment strength $r$ (x-axis) and food patch quality (y-axis). The variance is computed using Eq. (SE6). The system is simulated through SSA and we report the average of 1000 runs. The tests are for swarm size $S = 200$ and $n = 2$ food patches, with food patch 2's quality $q_2 = 0.5$ and food patch 1's quality varied on the y-axis in $[0.2, 1]$. In the full space the variance of the model with negative social feedback (right panel) is considerably lower than the variance of the other model's variance.

The mean-field model of the system with negative social feedbacks is described by the following ODE system:

$$\begin{aligned} \dot{x_i} &= q_i\, x_U + x_i(\rho\, q_i\, x_U - a - z\, x_i), \qquad i \in \{1, \dots, n\} \\ x_U &= 1 - \textstyle\sum_{i=1}^{n} x_i\ . \end{aligned} \tag{SE5}$$

We could not find the analytical solution for the system of Eq. (SE5) however numerical integration (*e.g.* see Figure 1 top-right of the main text) shows that the system can asymptotically converge to values very close to the target distribution of Eq. (1) of the main text. In all our simulations, through numerical optimisation we identified the best value of $z$ in terms of rapid convergence and small error (see more details in the Section *Parameters* below). Similarly to the system of Eq. (SE3), also the system of Eq. (SE5) has eigenvalues with different magnitude, however here the attraction is strong enough to move the system towards the fixed point in a finite time.

#### ST4. Variance reduction in the full space

We tested the two systems with a wide range of parameters in order to show that the results presented in the main text for specific values hold in a larger parameter space. Figure SF3 shows the variance of each model for varying the average recruitment strength $r$ (positive social feedback) and food patch quality. The reported variance is computed as

$$\text{Variance}(x_1 + x_2) = \text{Variance}(x_1) + \text{Variance}(x_2) + 2\,\text{Covariance}(x_1, x_2) \tag{SE6}$$

for the case of $n = 2$ food patches.

Figure SF4 shows the ratio between the variance of the system without negative feedback over the variance of the system with negative feedback. The ratio never goes below one and thus shows that using social negative feddback always reduces variance, in the tested systems.

Similarly, Figure SF5 shows how the variances are influenced by the swarm size $S$.

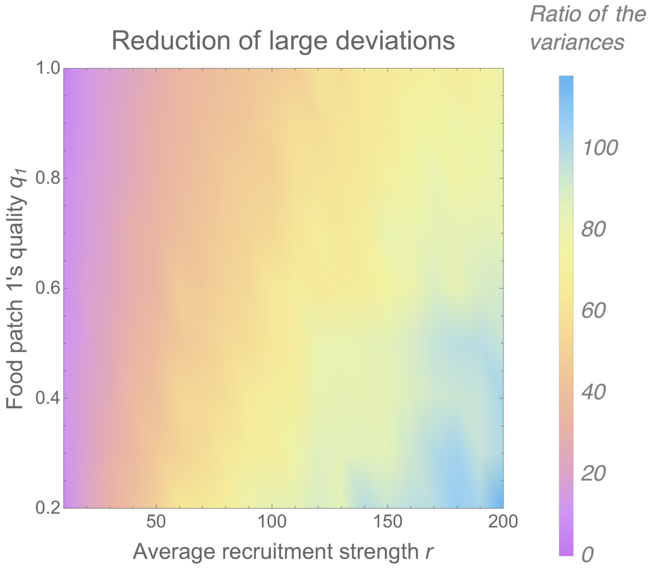


Figure SF4: Ratio of the variance of the system without and with negative social feedback (from Fig. SF3). The variance is computed using Eq. (SE6). The ratio never goes under one showing that the system with negative social feedback has always variance lower than the variance of the system without negative social feedback.

#### ST5. Effect of asocial negative feedback (abandonment or leak $\alpha$)

We tested the effect of increasing the negative feedback at the individual level (*i.e.* abandonment independent of social interaction) on the collective response. In the analyses reported in the main text and in the other supplementary texts we used a constant abandonment rate $\alpha = 10^{-3}$. Figure SF6 shows the stationary distribution of the subpopulations committed to the $n = 3$ food patches (same setup of Figure 1 of the main text) for different values of the abandonment rate $\alpha \in \{10^{-3}, 10^{-2}, 10^{-1}, 1, 10\}$ in the model without negative social feedback. While the ODE analysis predicts minimal deviations from the target distribution (Eq. (SE4)), the stochastic simulations evince high levels of variability for every tested value of $\alpha$. Therefore, our analysis suggests that, despite being a form of negative feedback, asocial leaking (abandonment) is not sufficient to guarantee low levels of variance in the collective distribution.

#### ST6. Adapting to changing conditions

When the starting point is different from the symmetric point $\{x_1, x_2, x_3\} = \{0, 0, 0\}$, the system without negative social feedback has very slow dynamics compared to the system with social negative feedback. An example is shown in Figure SF7 which shows the temporal evolution of the mean-field model for the starting point $\{x_1, x_2, x_3\} = \{0, 0.5, 0.5\}$. Such situations can happen when a change in the environment occur. While the system with negative social feedback reaches convergence with constant speed, unaffected by the starting point, the system without negative social feedback can only adapt in very long time.

#### ST7. Sum of squared error

We measure the performance of the system at convergence by computing the sum of squared error (SSE). The SSE is computed as

$$SSE = \sum_{i=1}^{n} \left( x_i^T - \frac{q_i}{\sum_{j \in n} q_j} \right)^2 . \tag{SE7}$$

where $x_i^T$ is the subpopulation committed to patch $i$ at convergence time $T = 1000$.

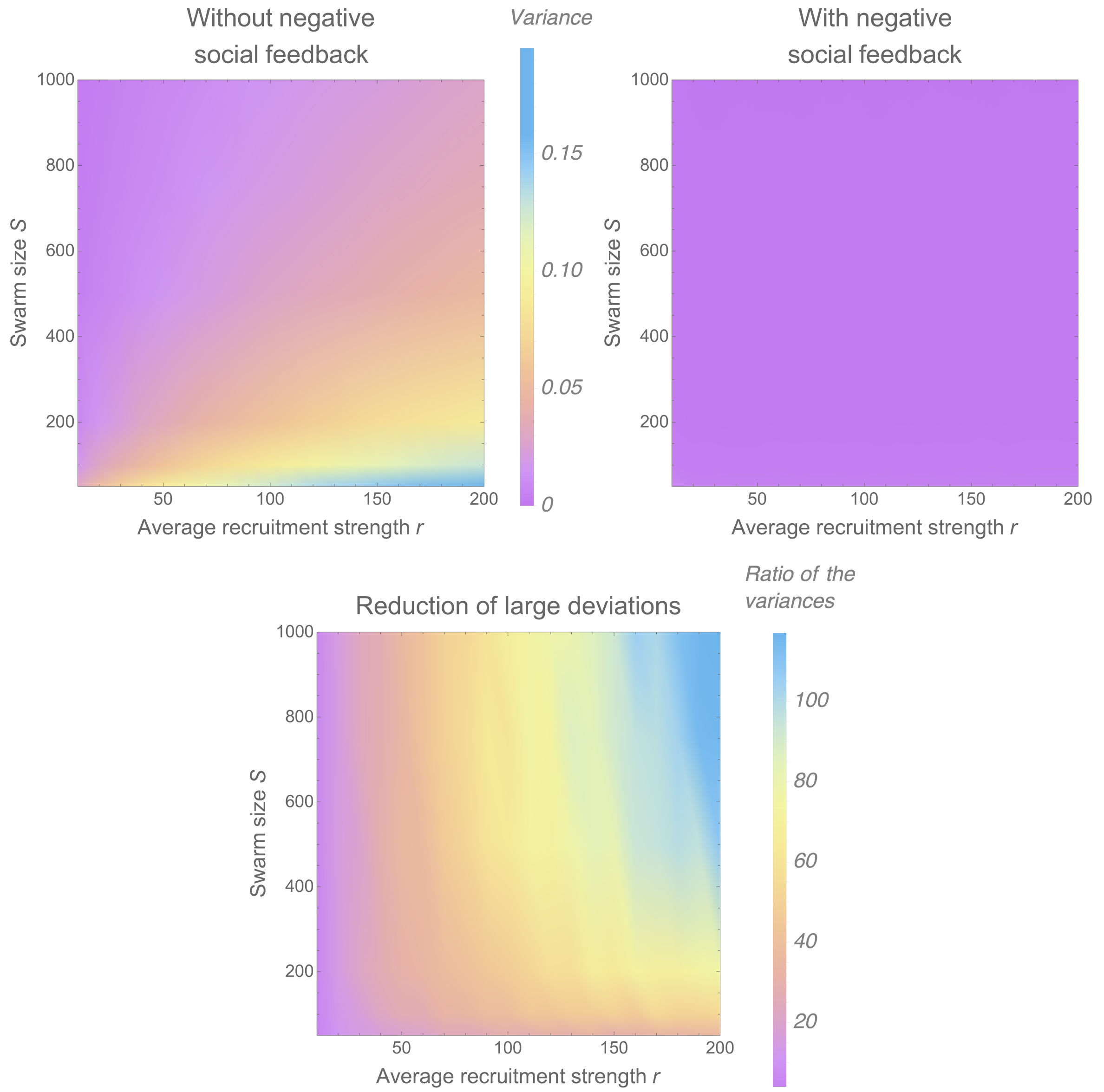


Figure SF5: Variance of the system for varying swarm size $S \in [50, 1000]$ and average recruitment strength $r \in [10, 200]$ for $q_1 = 1$ and $q_2 = 0.5$.. For any tested swarm size $S$, we observe the same pattern described in the main text.

Figure 2 of the main text shows how SSE—that is, the sum of the errors for each food source in achieving the target distribution—is significantly higher without negative social feedback.

#### ST8. Asocial model

By setting $\rho = 0$ in the model of Eq. (SE3), we remove any social component and the system only relies on individual discovery of the food patches and spontaneous abandonment. Figure SF8 shows that such a system has low variance centred around the target distribution. Additionally, the temporal dynamics, for the symmetric initial condition $\{x_1, x_2, x_3\} = \{0, 0, 0\}$, are comparable to the other two systems investigated in this study. The dynamics are slower when the starting point is different from $\{x_1, x_2, x_3\} = \{0, 0, 0\}$. Figure SF9 illustrates an example for the starting point $\{x_1, x_2, x_3\} = \{0, 0.5, 0.5\}$. We note, however, that the system can speed up its dynamics by increasing the leak rate (*i.e.* spontaneous abandonment $a$) as illustrated in the right panel of Figure SF9. The

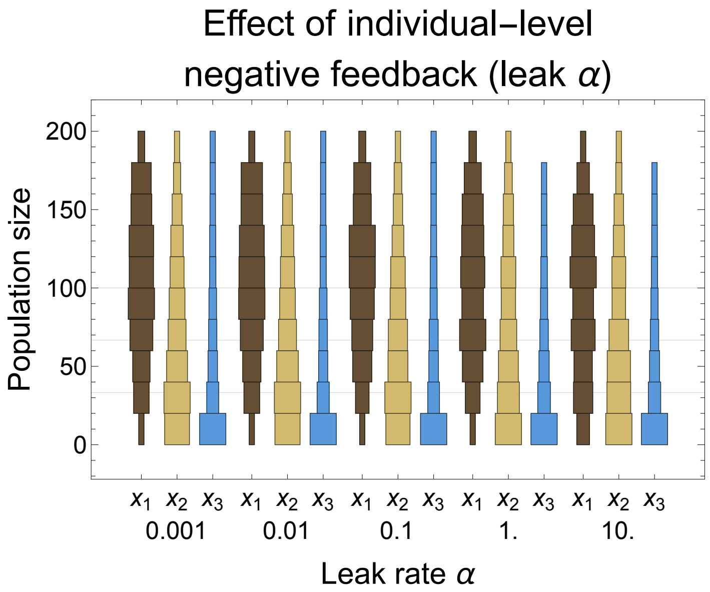


Figure SF6: Results from $10^3$ SSA simulations of the model without negative social feedback for different values of the abandonment rate $\alpha$ (leak), changed on the x-axis. We ran simulations of swarms composed of $S = 200$ individuals operating in an environment with $n = 3$ food patches, with the same quality values used for Figure 1 of the main text, $(q_1 = 0.75, q_2 = 0.5, q_3 = 0.25)$, and the average recruitment strength is $r = 100$. The distribution charts display how the population is divided, at the final step (at time 10), among the three sub-populations foraging from the different patches $(x_1, x_2, x_3)$, labelled on the axis and represented with three different colours. Variance remains high also for high levels of independent (asocial) negative feedback.

spontaneous abandonment can be increased however not unboundedly as the parameter $a$ impacts on how close to the target distribution the system converges to. In fact, the predicted fixed point for $\rho = 0$ in the model of Eq. (SE3) is:

$$x_i^* = \frac{q_1}{a + q_1 + q_2} \,. \tag{SE8}$$

#### ST9. Large deviation from target

Figure SF11 shows the probability that the state of the system at convergence would deviate from the target. We set an acceptance margin $m$ and we measure the probability of laying outside such a margin. The results show the proportion of 1000 SSA runs where at least one of the subpopulations committed to either of the $n = 2$ food patches has a deviation larger than $m\,S$ (where $m$ is the given margin and $S$ the system size). In Figure SF11, we vary the margin value $m$ and show that the probability of being outside any given margin is higher without negative social feedback.

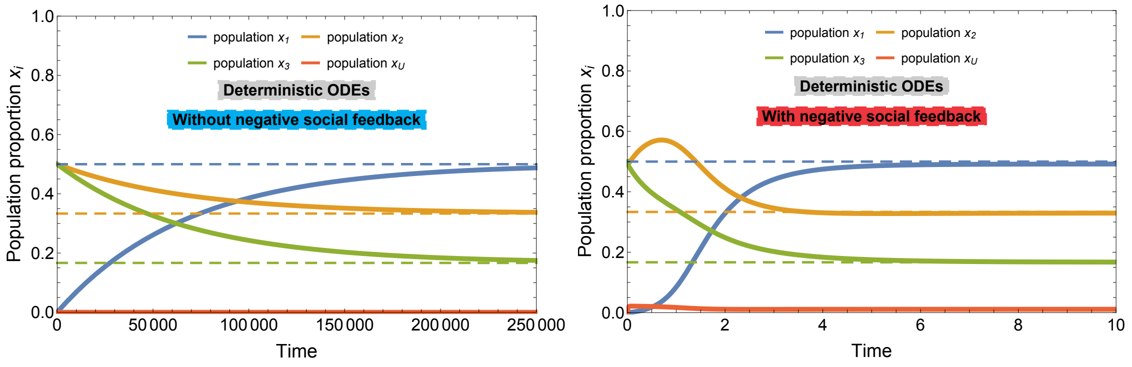


Figure SF7: ODE dynamics over time for starting point $\{x_1, x_2, x_3\} = \{0, 0.5, 0.5\}$. The system with negative social feedback (right) converges to the target distribution in a similar amount of time for every starting point. Instead, the system without negative social feedback (left) has very slow dynamics and the time to converge to the target distribution is more than five orders of magnitude larger than the symmetric case presented in the main text (Figure 1).

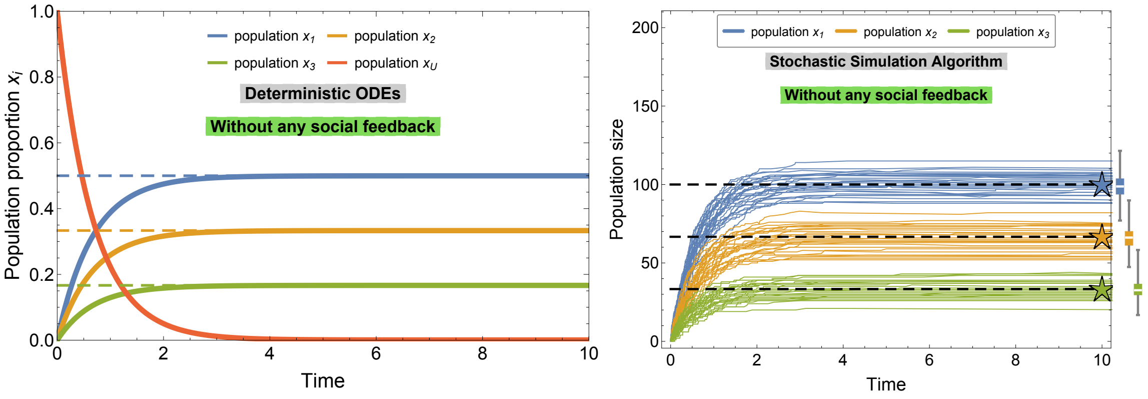


Figure SF8: Dynamics of the ODE model (left) and SSA simulations (right) of the system without social feedback with starting point $\{x_1, x_2, x_3\} = \{0, 0, 0\}$. The system without social feedback also displays small variance (boxplots on the right for 1000 simulations), however the dynamics can be slow for different starting points, *e.g.* see Figure SF9.

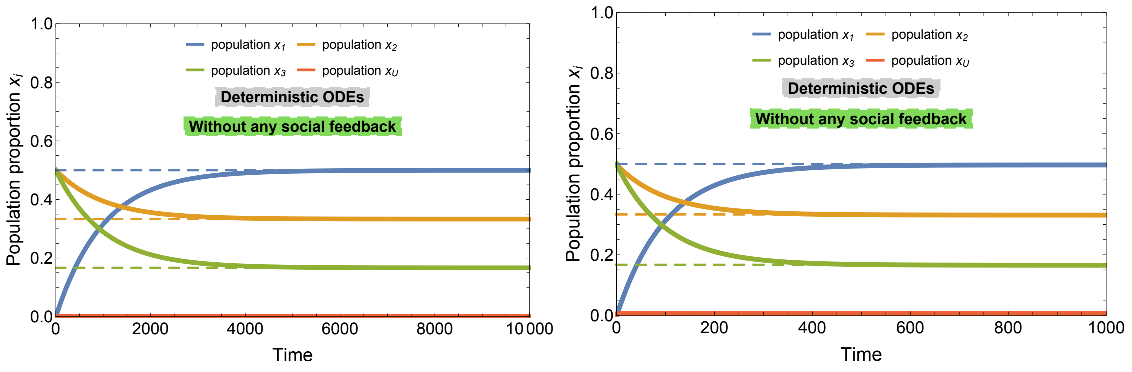


Figure SF9: ODE dynamics of the system without social feedback with starting point $\{x_1, x_2, x_3\} = \{0, 0.5, 0.5\}$. In both cases, the convergence time is longer than the symmetric condition of Figure SF8. On the left panel, the leak rate $a = 10^{-3}$ is the same of the other analysed models. On the right panel, we show that the system can speed up the dynamics by increasing the leak rate to $a = 10^{-2}$.

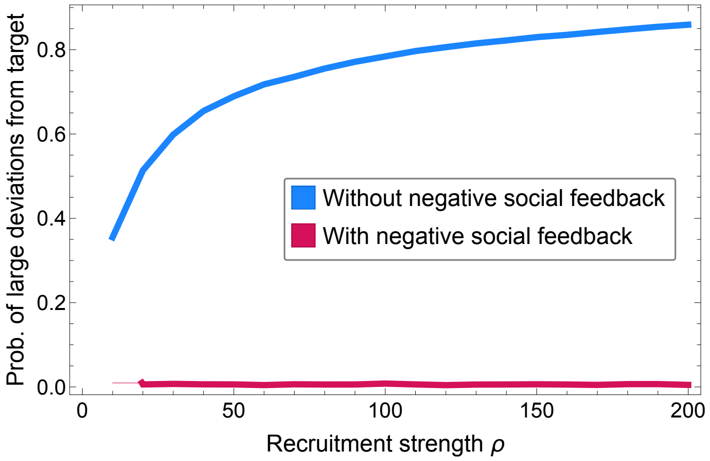


Figure SF10: Probability of large deviations from the target distribution for a system of $S = 200$ individuals computed from $10^3$ SSA simulations. The probability is computed as the percentage of simulations that have the final population distribution with a deviation larger than $10\%S$ from the target distribution. The results are an average for $q_1 \in [0.2, 1]$ and $q_2 = 0.5$. The system with negative feedback shows a large reduction of the probability of large deviations. (95% confidence intervals are smaller of the line width.)

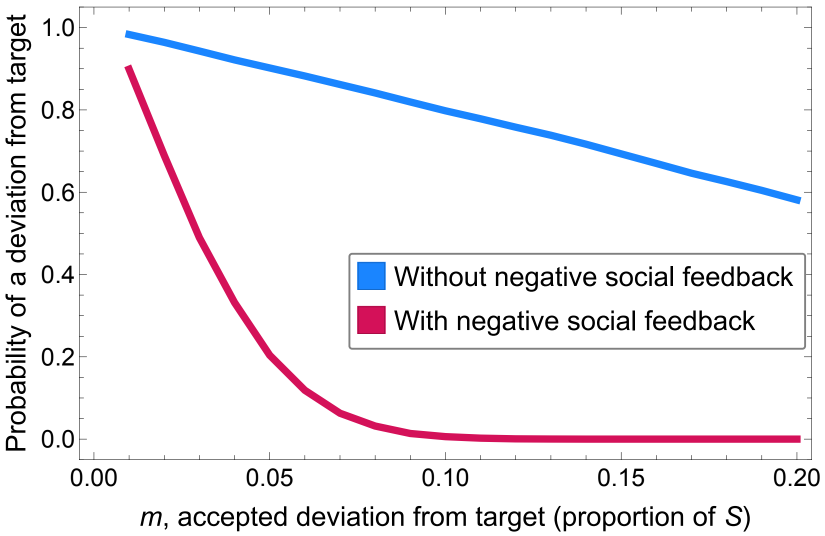


Figure SF11: We compute the probability of a deviation from the target distribution larger than a threshold $m$, where $m$ is varied on the x-axis and the swarm size is $S = 200$. We report here the proportion of $10^3$ SSA runs that had a deviation larger than $m$ (y-axis) for an average recruitment strength $r = 100$. The system with negative social feedback has always a lower probability than the system without negative social feedback. (95% confidence intervals are smaller of the line width.)
